# Transcriptome profiling reveals bisphenol A alternatives activate estrogen receptor alpha in human breast cancer cells

**DOI:** 10.1101/112862

**Authors:** Robin Mesnage, Alexia Phedonos, Matthew Arno, Sucharitha Balu, J. Christopher Corton, Michael N Antoniou

## Abstract

**Background:** Plasticizers with estrogenic activity, such as bisphenol A (BPA), have been reported to have potential adverse health effects in humans. Due to mounting evidence of these health effects and public pressure, BPA is being phased out by the plastics manufacturing industry and replaced by other bisphenol variants in “BPA-free” products.

**Objectives:** We have compared estrogenic activity of BPA to 6 bisphenol analogues (bisphenol S, BPS; bisphenol F, BPF; bisphenol AP, BPAP; bisphenol AF, BPAF; bisphenol Z, BPZ; bisphenol B, BPB) in three human breast cancer cell lines.

**Methods:** Estrogenicity was assessed by cell growth in an estrogen receptor (ER)-mediated cell proliferation assay, and by the induction of estrogen response element (ERE)-mediated transcription in a luciferase assay. Gene expression profiles were determined in MCF-7 human breast cancer cells by microarray analysis and confirmed by Illumina-based RNA sequencing.

**Results:** All bisphenols showed estrogenic activity in promoting cell growth and inducing ERE-mediated transcription. BPAF was the most potent bisphenol, followed by BPB > BPZ ~ BPA > BPF ~ BPAP > BPS. The addition of ICI 182,780 antagonized the activation of ERs by bisphenols. Data mining of ToxCast high-throughput screening assays confirms our results but also shows divergence in the sensitivities of the assays. The comparison of transcriptome profile alterations resulting from BPA alternatives with an ERα gene expression biomarker further indicates that all BPA alternatives act as ERα agonists in MCF-7 cells. These results were confirmed by RNA sequencing.

**Conclusion:** In conclusion, BPA alternatives are not necessarily less estrogenic in a human breast cancer cell model. Three bisphenols (BPAF, BPB, and BPZ) were more estrogenic than BPA. The relevance of human exposure to BPA alternatives in hormone-dependent breast cancer risk should be investigated.

## Introduction

Plasticizers such as bisphenol A (BPA) have been reported to have potential adverse health effects in humans, including reproductive endocrine disorders and neurobehavioral problems (Rochester, 2013). BPA is one of the best-studied endocrine disrupting chemicals (EDCs), with more than 75 out of 91 published studies showing associations between BPA exposure and human health as of May 2013 (Rochester, 2013). The primary mode of action of BPA is the activation of estrogen receptor (ER) mediated transcription (Shioda *et al.,* 2013). Biomonitoring studies suggest that the urine from a majority of individuals in industrialized countries contain measurable BPA and metabolites (Gerona *et al.,* 2016). BPA is detected in breast milk (Nakao *et al.,* 2015). A causal link between BPA and breast cancer remains equivocal because epidemiological studies have reported conflicting results (Rochester, 2013). Nonetheless, there is ample evidence from laboratory rodent and non-human primate studies that prenatal exposure to BPA may increase the propensity to develop breast cancer during adulthood (Mandrup *et al.,* 2016; Tharp *et al.,* 2012). In these studies, low-doses of BPA has been shown to affect mammary gland development by causing hyperplastic lesions in rats, which are parallel to early signs of breast neoplasia in women (Mandrup et al., 2016). These findings suggests that mammary tumors provoked by BPA may be initiated in the womb, and thus more prospective studies measuring BPA in utero are needed in order to conclude there is a carcinogenic potential for humans. Due to mounting evidence of harm and public pressure, BPA is being phased out by plastics manufacturers and is being replaced by other BP variants in “BPA-free” products (Rochester and Bolden, 2015).

Bisphenol F (BPF), Bisphenol S (BPS), and Bisphenol AF (BPAF) are among the main substitutes of BPA in polycarbonate plastics and epoxy resins (Chen *et al.,* 2016). They are commonly found in baby feeding bottles, on thermal receipt papers, glues, dental sealants, food packaging or personal care products. There is a strong negative correlation between the levels of BPA and BPS on thermal paper, whereby the papers containing high quantities of BPS have low quantities of BPA, suggesting that BPS has in part replaced BPA in this context (Liao *et al.,* 2012). The handling of a thermal receipt paper before eating French fries is sufficient to cause a rapid increase of serum BPA levels within 30 min (Hormann *et al.,* 2014). Epidemiological data on the human body burden of different BPA alternatives is very limited, but evidence suggests that they are generally widespread. In an investigation of the presence of BPS and bisphenol F (BPF) in human urine, these compounds were found to be present in 78% and 55% of respective samples in the United States (Zhou *et al.,* 2014). Another investigation revealed that although the average urinary concentration of BPA in 130 individuals from Saudi Arabia was 4.92 ng/mL, 7 other bisphenols were found with the total bisphenol concentration reaching 19 ng/mL (Asimakopoulos *et al.,* 2016), suggesting that BPA is only one of many BP alternatives that are found in humans.

Some of these BPA replacements are structurally related to BPA and have endocrine disrupting effects. Numerous studies have suggested that BPS and BPF have potencies similar to that of BPA resulting in endocrine disrupting effects such as uterine growth in rodents (Rochester and Bolden, 2015). A number of in vitro studies have looked at endocrine mode of action of some BPA alternatives in nuclear hormone receptor interactions (Chen et al., 2016). The range of EC50 values for estrogen activity (0.02 μM to >1000 μM) and androgen activity (0.29 μM to >1000 μM) of 20 BP congeners (5 of which are being tested here) indicates remarkable differences (Kitamura *et al.,* 2005). Another study showed that 5 BPA alternatives (3 studied here) exhibited potencies within the same range as BPA on steroidogenesis, androgen receptor, aryl hydrocarbon receptor, and retinoic acid receptor activity (Rosenmai *et al.,* 2014). Some BPA alternatives have also been tested in ER high-throughput screening assays in the context of the US Environmental Protection Agency (EPA) ToxCast program and shown in some cases to have estrogen-like effects (Judson *et al.,* 2015).

Perturbation of the gene expression profile of a cell by chemicals results in unique signatures, allowing comparisons between transcriptome alterations caused by drugs and diseases to predict therapeutic and off-target effects (Iorio *et al.,* 2010). The clinical utility of gene expression profiles is exemplified by studies showing that transcriptome signatures may affect clinical decision-making in breast cancer (Rhodes and Chinnaiyan, 2005). Gene expression have been shown to be a more powerful predictor of breast cancer survival than standard systems based on clinical and histologic criteria (van de Vijver *et al.,* 2002). Exposure of breast cancer cells to BPA present a gene expression profile of tumor aggressiveness associated with poor clinical outcomes for breast cancer patients (Dairkee *et al.,* 2008). While ER activation drives two-thirds of breast cancer (Green and Carroll, 2007), there have been no studies aimed at elucidating transcriptome alterations underlying estrogenic effects of a broad range of BPA alternatives to which human populations are exposed. In order to address this deficiency, we studied alterations in gene expression profiles of estrogen-dependent MCF-7 human breast cancer cells caused by BPA and 6 alternatives (Figure 1) using Affymetrix microarray and Illumina RNA sequencing platforms. These BPA alternatives are detected in different categories of food items (Liao and Kannan, 2013) and are found in human biological fluids (Asimakopoulos et al., 2016). The transcriptome signatures obtained were compared to a gene expression biomarker, which has been demonstrated to accurately predict ER modulation, including by “very weak” agonists (Ryan *et al.,* 2016). We also used an E-screen assay (cell proliferation in estrogen-responding cells) as well as an estrogen response element (ERE)-luciferase reporter gene assay system. Our results clearly demonstrate that all 6 BPA alternative compounds are estrogenic. Alterations of MCF-7 transcriptome profiles caused by BPA alternatives bear the signature of ER activation with three bisphenols (BPAF, BPB, BPZ) having more estrogenic potency than BPA.

**Figure 1.**
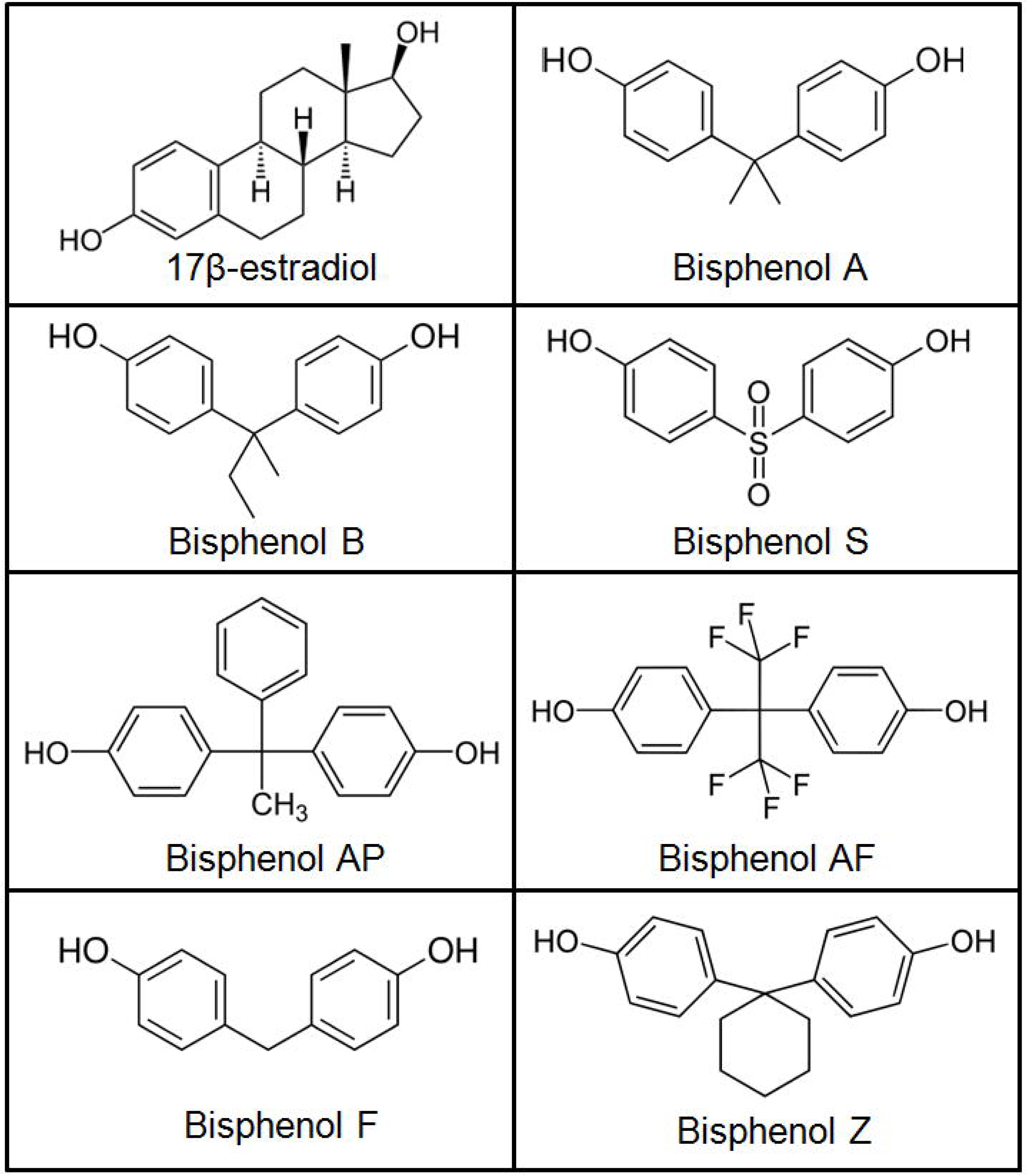
Molecular structures of BPA and bisphenol alternatives in comparison to the natural hormone 17 β-estradiol.

## Methods

### Cell culture and treatments

All reagents and chemicals, unless otherwise specified, were of analytical grade and purchased from Sigma-Aldrich (UK). MCF-7, MDA-MB-231 and T47D human breast cancer cell lines were a gift from Prof. Joy Burchell (Research Oncology Department, King’s College London, UK). The T47D-KBluc cells were purchased from the American Type Culture Collection (ATCC, Teddington, UK) and harbour a stably integrated copy of a luciferase reporter gene under control of a promoter containing EREs. All cells were grown at 37°C (5% CO_2_) in 75 cm^2^ flasks (Corning, Tewksbury, USA) in a maintenance medium composed of phenol red free DMEM (Life Technologies, Warrington, UK), 10% fetal bovine serum (FBS; GE Healthcare Life Sciences, Buckinghamshire, UK), 2mM glutamine (GE Healthcare Life Sciences) and 10 μg/ml penicillin/streptomycin (Life Technologies). Test substances were diluted in 100% ethanol to prepare stock solutions. All treatments were prepared in a test medium containing phenol red free DMEM, 5% charcoal stripped FBS (Life Technologies), 2mM glutamine (GE Healthcare Life Sciences) and 10 μg/ml penicillin/streptomycin (Life Technologies). The solvent control contained 0.1% (v/v) absolute ethanol diluted in test medium. Cells were detached from the flask substrate using 0.05% trypsin EDTA (Life Technologies) and counted using a haemocytometer prior to seeding. After a 24 h recovery period in DMEM maintenance medium tests were applied.

### E-Screen assay

The E-screen assay allows the determination of estrogenic effects by determining estrogen receptor-mediated cell proliferation in hormone-responsive cells. This assay was performed as originally described (Soto *et al.,* 1995), except that the bioassay was terminated using a MTT assay, indirectly measuring cell number by testing the activity of mitochondrial succinate dehydrogenase, in order to assess cell proliferation. Briefly, cells were seeded into 48-well plates (Dutscher Scientific, Brentwood, UK) at a density of 20,000 cells per well in 250 μl maintenance medium. Following a 24-hour incubation to allow cell attachment, medium was changed to include various concentrations of treatment substances. The test medium was refreshed after 3 days. After another 3 day incubation, an MTT assay was performed as follows. Cells were incubated with 250 μl of MTT solution (1 mg/ml) for 2 hours and the test terminated by lysing the cells with dimethyl sulfoxide (DMSO). The optical density of the cell lysate was measured at 570 nm using the SPECTROstar Nano plate reader (BMG Labtech, Ortenberg, Germany). The cell proliferative effect was expressed as a percentage of the control, untreated samples.

## ERE-mediated luciferase reporter gene assay

The ERE-mediated transcription of a luciferase reporter gene (3×ERE) was determined in the T47D-KBluc cells (Wilson *et al.,* 2004)using the Steady-Glo^®^ luciferase assay system following the manufacturer’s instructions (Promega, Southampton, UK). The T47D-KBluc cells were seeded in white 96 well plates (Greiner Bio-One, Germany) at a density of 20,000 cells per well in 50 μl of maintenance medium and allowed to attach overnight. An initial 24 h incubation was performed in the absence of test substances to purge cells of hormone residues in order to improve estrogen deprivation. Test substances were added and plates were incubated for another 24 hours before the addition of 50 μl Steady-Glo^®^ luciferase reagent. The plates were left to stand for 10 minutes in the dark at room temperature to allow cell lysis and establishment of the luciferase reaction. Luminescence was measured using the Orion II microplate luminometer (Berthold Detection Systems, Bad Wildbad, Germany). Estrogen receptor modulation was confirmed by measuring the inhibitory effect of ICI 182,780 on the concentration of BPA resulting in a 20-fold increase in luciferase activity.

## Microarray gene expression profiling

MCF-7 cells were seeded into 96-well plates with maintenance medium at a density of 20,000 cells per well. After 24 h of steroid hormone deprivation in hormone-free medium, the cells were stimulated with test substances for 48 h in triplicate in three independent experiments. RNA extraction was performed using the Agencourt RNAdvance Cell V2 kit according to the manufacturer's instructions (Beckman Coulter Ltd, High Wycombe, UK). The samples were checked for RNA quality using the Agilent 2100 Bioanalyzer (Agilent Technologies LDA UK Limited, Stockport, UK) and quantified using the Nanodrop ND-1000 Spectrophotometer (Thermo Fisher Scientific, Wilmington, USA). The RNA integrity number (RIN) was 9.7 ± 0.3. Subsequently, triplicate samples which passed quality control (QC) criteria were pooled appropriately such that the final input amount of each sample was 3 ng.

The gene expression profiles were determined using the Affymetrix Human Transcriptome 2.0 Array as follows. Single Primer Isothermal Amplified (SPIA) cDNA was generated using the Ovation Pico WTA System V2 kit (Nugen, AC Leek, The Netherlands) following the manufacturer's instructions. In addition, the SPIA cDNA was subjected to a QC check to assess quality (Agilent 2100 Bioanalyzer) and quantity (Nanodrop ND-1000 Spectrophotometer) in preparation for the next stage. The SPIA cDNA was fragmented and biotin-labelled using the Encore Biotin Module (Nugen) according to the manufacturer’s instructions. The fragmented and biotin-labelled cDNA was subjected to a further round of QC checks to assess fragmentation size (Agilent 2100 Bioanalyzer). Hybridisation cocktails were prepared from the fragmented labelled-cDNA according to Nugen's recommendations and hybridised to the microarrays at 45°C overnight. The arrays were washed and stained using the wash protocol FS450_0001 recommended for Affymetrix Human Transcriptome 2.0 Arrays on the GeneChip Fluidics 450 station. Ultimately, the arrays were scanned using the Affymetrix GeneChip Scanner. CEL files were QC assessed in the Expression Console software package (Affymetrix) by using standard metrics and guidelines for the Affymetrix microarray system. Data was imported and normalised together in Omics Explorer 3.0 (Qlucore, New York, NY, USA), using the Robust Multi-array Average (RMA) sketch algorithm. These microarray data have been submitted to Gene Omnibus and are accessible through accession number GSE85350.

## Gene expression profiling by RNA-sequencing

RNA-sequencing was performed by applying Illumina sequencing by synthesis technology as follows. The amount of RNA for each library (100 ng) was a pool made up of 33 ng of RNA from each of the replicate wells for each sample. The preparation of the library was done by NEBNext Ultra Directional RNA (New England Biolabs, Hitchin, UK) following the manufacturer’s protocol. The amplified library was assessed using the Agilent 2100 Bioanalyzer for size and presence of adapter/primers/dimers – sized at ~400bp (including ~130bp adapter). The rRNAs were removed using the rRNA depletion module (New England Biolabs) following the manufacturer’s protocol. Libraries were pooled together and sequenced on a HiSeq2500 using a Rapid Run v2 flowcell with on-board clustering in a 2x100 paired-end (PE) configuration. BCL files were processed and deconvoluted using standard techniques. The sequencing output FASTQ files contain the sequences for each read and also a quality score. We have analysed the quality scores and other metrics using FASTQC (http://www.bioinformatics.babraham.ac.uk/projects/fastqc/). Contamination from rRNA was measured using an alignment script (http://genomespot.blogspot.co.uk/2015/08/screen-for-rrna-contamination-in-rna.html). Adapter sequences (standard TruSeq LT adapter seq) were removed/trimmed using cutadapt (Martin, 2011). Sequences were then aligned to the genome (hg38 database) using the hierarchical indexing for spliced alignment of transcripts program HISAT2 (Kim *et al.,* 2015). BAM files were imported to Qlucore omics explorer, along with the GTF file for known genes in hg38 (downloaded from UCSC). Qlucore normalises data using a method similar to the trimmed mean of M-values normalization method (TMM), which corrects for transcript length and applies a log transformation (Robinson and Oshlack, 2010) and gives values similar to the quantification of transcript levels in reads per kilobase of exon model per million mapped reads (RPKM) (Robinson and Oshlack, 2010), but also incorporates the TMM normalization factor for an improved between-sample normalization.

A total of 376.6 million raw reads were obtained (25.1 ± 5.4 million reads per sample). The average value of Q30, representing the probability of an incorrect base call 1 in 1,000 times, was above 96%. The GC content (%GC) of the reads was on average 49%. A total of 90.92 ± 6.54% of the clean reads were mapped onto the human reference genome hg38. Among them, an average of 71.01 ± 5.82 % and 18.25 ± 3.79 % reads align concordantly exactly one time and more than one time, respectively. These RNA-seq data have been submitted to Gene Omnibus and are accessible through accession number GSE87701.

## ToxCast data mining

We analyzed publically available data from the ToxCast program using the iCSS ToxCast Dashboard (http://actor.epa.gov/dashboard/). In the ToxCast program, out of 18 ER assays, 5 measured ERE-mediated transcription and gave results comparable to those of our T47D-Kluc assay. One assay was a single-readout fluorescent protein induction system measuring interaction of ER with ERE at two time points by microscopy technology (OT_ERa_EREGFP_0120 and _0048). Two other assays were reporter gene assays measuring mRNA in HepG2 cells (ATG_ERa_TRANS_up and ATG_ERE_CIS_up). The last two assays measured reporter protein levels in HEK293T (To×21_ERa_BLA_Agonist_ratio) and BG-1 (To×21_ERa_LUC_BG1_Agonist) cell lines. Other assays of protein complementation that measure formation of ER dimers and test for activity against ERα (OT_ER_ERaERa_0480 and _1440). Finally, NVS_NR_bER and NVS_NR_hER are radioligand receptor binding assays. Not all assays were performed on all bisphenols. For instance, BPAP was only tested in Attagene assays. In this study, the AC50 value was used as a quantitative measure to reflect the potency of BP alternatives.

## Comparison of the microarray and RNA-Seq data to the ER biomarker

A rank-based nonparametric analysis strategy called the Running Fisher test implemented within the NextBio database (Illumina, San Diego, CA) was used to compare gene lists derived from the microarray and RNA-Seq data to the ER biomarker characterized earlier (Ryan et al, 2016). The Running Fisher test is a normalized ranking approach which enables comparability across data from different studies, platforms, and analysis methods by removing dependence on absolute values of fold-change, and minimizing some of the effects of normalization methods used, while accounting for the level of genome coverage by the different platforms. The Running Fisher algorithm computes statistical significance of similarity between ranked fold-change values of two gene lists using a Fisher exact test (Kupershmidt *et al.,* 2010). A p-value was selected as our cutoff for significance based on prior evaluation of the cutoff as predictive of activation of ER (Ryan et al., 2016).

## Statistical analysis

The concentration required to elicit a 50% response (AC50) was determined using a nonlinear regression fit using a sigmoid (5-parameters) equation calculated with GraphPad software (GraphPad Software, Inc., La Jolla, CA, USA). For the transcriptome analysis, pairwise comparisons of each tested substance to the negative control were performed using a t-test controlling for batch effects in Omics Explorer 3.0 (Qlucore). Data used for the functional analysis were selected at cut-off p-values < 0.05 with fold-change > 1.2 to study the ER activation signature as previously described (Ryan et al., 2016). Gene and disease ontology were analyzed using the Thomson Reuters MetaCore Analytical Suite version 6.28 recognizing network objects (proteins, protein complexes or groups, peptides, RNA species, compounds among others). The p-values are determined by hypergeometric calculation and adjusted using a Benjamini & Hochberg approach. All experiments were performed 3 times in triplicate (n=3).

## Results

### E-screen and ERE-luciferase reporter gene assays

We have studied the estrogenic activities of 7 BP found in foodstuffs and human biological fluids in three human breast cancer cell lines. As an initial investigative step of estrogenic potential, an E-Screen assay was performed using two hormone dependent (MCF-7, T47D) and one hormone independent (MDA-MB-231) human breast cancer cell lines (Figure 2A). The positive control 17β-estradiol was very potent in inducing the proliferation of MCF-7 cells (AC50 = 8 pM). All BP derivatives were also able to promote cell growth at concentrations 10,000 to 100,000 higher than 17β-estradiol (Figure 2A, top panel). BPAF was the most potent BP (AC50=0.03 μM) followed by BPZ (0.11 μM) > BPB (0.24μM) > BPA (0.36 μM) > BPAP (0.39 μM) > BPF (0.55 μM) > BPS (1.33 μM). The same overall trend was observed with the T47-D cell line (Figure 2A, middle panel), albeit to a lesser extent which can be accounted for by the fact that T47-D cells express lower levels of ER. As expected no cell proliferative effects were observed with the hormone independent, ER-negative MDA-MB-231 cell line suggesting that proliferative effects were mediated by ER (Figure 2A, bottom panel).

**Figure 2.**
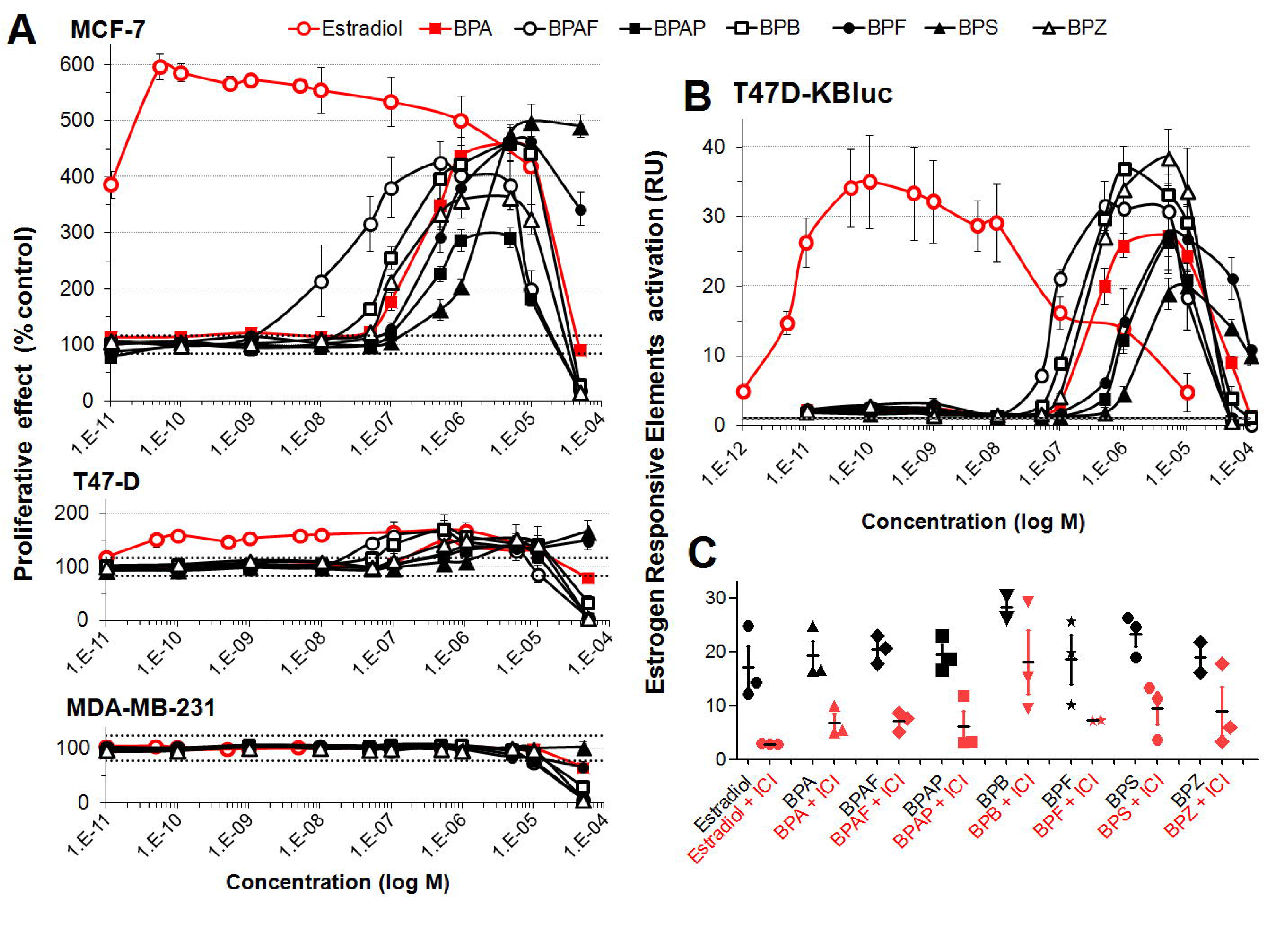
Bisphenol A alternatives can effectively substitute for estradiol in promoting growth through estrogen receptors of human breast cancer cells. **(A)** Proliferative effect of BPA and bisphenol alternatives on mammary cells in an E-Screen bioassay. After 24h of steroid hormone starvation cells were treated for 6 days with the test compounds. Numbers of cells was measured by an MTT colorimetric assay. Results are expressed as proliferative effect percentage relative to the proliferation of cells under hormone-free conditions. Data are the mean +/- SE of three independent experiments, with each performed in triplicate. **(B)** Bisphenol A alternatives stimulate estrogen response element-mediated transcription. After 24 hours of steroid hormone starvation, T47D-Kluc cells were treated with the test compounds for 24 hours. Cells were then lysed and subjected to a bioluminescence luciferase reporter gene assay. **(C)** Luciferase assays of T47D-Kluc cells treated with bisphenol A alternatives in the absence (black) or presence (red) of ICI 182,780, an estrogen receptor antagonist. Results show that ICI 182,780 represses ERE-mediated transcription induced by bisphenol A alternatives.

Since the observed BP-derivative induced proliferation of MCF-7 and T47D cells (Figure 2A and 2B) could be mediated by different hormones, estrogenic effects of BPA alternatives was then investigated employing a luciferase reporter gene assay allowing measurement of ERE-mediated transcription. The results (Figure 2B, upper panel) were very similar to those obtained from the E-Screen assay: BPAF was the most potent bisphenol (AC50 = 0.08 μM) at stimulating ERE-luciferase reporter gene expression followed by BPB (0.3 μM) > BPZ (0.4 μM) ~ BPA (0.4 μM) > BPF (1 μM) ~ BPAP (1 μM) > BPS (1.5 μM) having estrogenic effects in the same range as BPA. Moreover, the addition of ICI 182,780 (100 nM) antagonized the activation of ER by estradiol and bisphenols confirming reporter gene expression via this receptor (Figure 2C). However, estrogenic effects of some BP derivatives, such as BPB and BPZ, were not completely antagonized by 100 nM ICI addition, suggesting estrogenic activation mechanisms independent of ER.

ToxCast data mining (Table 1) confirms our results but also reveals some discrepancy with the results of the ToxCast assays. BPF, BPZ and BPB were classified as inactive in the ToxCast gene reporter ERa_BLA_Agonist_ratio assay, because they did not reach the response threshold, possibly due to the lower sensitivity of this assay. BPF was classified as inactive in the majority of ToxCast assays although it has been found to be positive in the EDSP Tier 1 uterotrophic assay (Judson et al., 2015).

**Table 1.** Comparison to ToxCast data. Publically available data was analysed from the ToxCast program using the iCSS ToxCast Dashboard. AC50 values (μM) for ToxCast assays relative to ESR1 activation are compared to the results presented here. Negative results are displayed by grey boxes. White boxes shows assays in which bisphenols have not been tested.

| CASRN | Name | ATG_ERa_TRANS_up | ATG_ERE_CIS_up | NVS_NR_bER | NVS_NR_hER | OT_ER_ERaERa_0480 | OT_ER_ERaERa_1440 | OT_ERa_EREGFP_0120 | OT_ERa_EREGFP_0480 | TOX21_ERa_BLA_Agonist_ratio | TOX21_ERa_LUC_BG1_Agonist | AC50 MCF7 this study | AC50 LUC this study |
| --- | --- | --- | --- | --- | --- | --- | --- | --- | --- | --- | --- | --- | --- |
| 1478-61-1 | Bisphenol AF | 0.36 | 0.09 | 0.07 | 0.03 | 1.05 | 0.712 | 0.1 | 0.11 | 0.36 | 0.05 | 0.03 | 0.08 |
| 843-55-0 | Bisphenol Z |  |  |  |  |  |  |  |  | #### | 0.31 | 0.11 | 0.4 |
| 77-40-7 | Bisphenol B | 0.13 | 0.17 | 0.21 | 0.27 | 1.82 | 1.41 | 0.18 | 0.22 | #### | 0.11 | 0.24 | 0.3 |
| 80-05-7 | Bisphenol A | 0.12 | 0.1 | 0.42 | 0.23 | 5.73 | 4.33 | 0.42 | 0.63 | 1.01 | 0.36 | 0.36 | 0.4 |
| 1571-75-1 | Bisphenol AP | 0.12 | 0.29 |  |  |  |  |  |  |  |  | 0.39 | 1 |
| 2467-02-9 | Bisphenol F | 34.9 | 30.3 | #### | #### | #### | #### | #### | #### | #### | #### | 0.55 | 1 |
| 80-09-1 | Bisphenol S | 0.8 | 1.42 | 17.7 | 5.45 | 43.8 | 56.1 | 59.2 | 26.6 | 9.29 | 1.87 | 1.33 | 1.5 |

### Transcriptome profiling

MCF-7 cells treated for a period of 48h with BPA, 6 BP alternatives, or 17β-estradiol, were then subjected to full transcriptome profiling using the Affymetrix microarray platform. The length of exposure (48h) and the concentration of test agent were selected, because previous studies showed that these conditions produce robust gene expression changes (Shioda *et al.,* 2006). Supplementary File 1 shows the statistical significance of differential transcript cluster expression with respective fold-changes by volcano plots. The set of genes altered by treatment with BPA and BP alternatives (Supplementary File 2) were highly enriched in genes involved in the regulation of the cell cycle (Figure 4A), as well as in genes regulated by steroid hormones (Figure 4B). Additionally, an enrichment analysis of MeSH terms reveals that exposure to both BPA or the 6 BP alternatives are significantly enriched in genes involved in the etiology of breast cancer (Figure 4C) such as *ALDH1A3, CAMK1D, PPARG* or *PGR.* Genes having the highest fold-changes were similar and indicative of a hormone-induced proliferative effect (Table 2). The gene encoding the progesterone receptor *(PGR)* consistently exhibited the highest fold-change after stimulation with BPA or all the BP alternatives. ER binding motifs were overrepresented in the promoter of the differentially expressed genes (DEGs). There were 4.8 to 6.4 more ER binding sequences (EREs) in the promoters of the DEGs than what would be expected by chance (Figure 3D).

**Figure 3.**
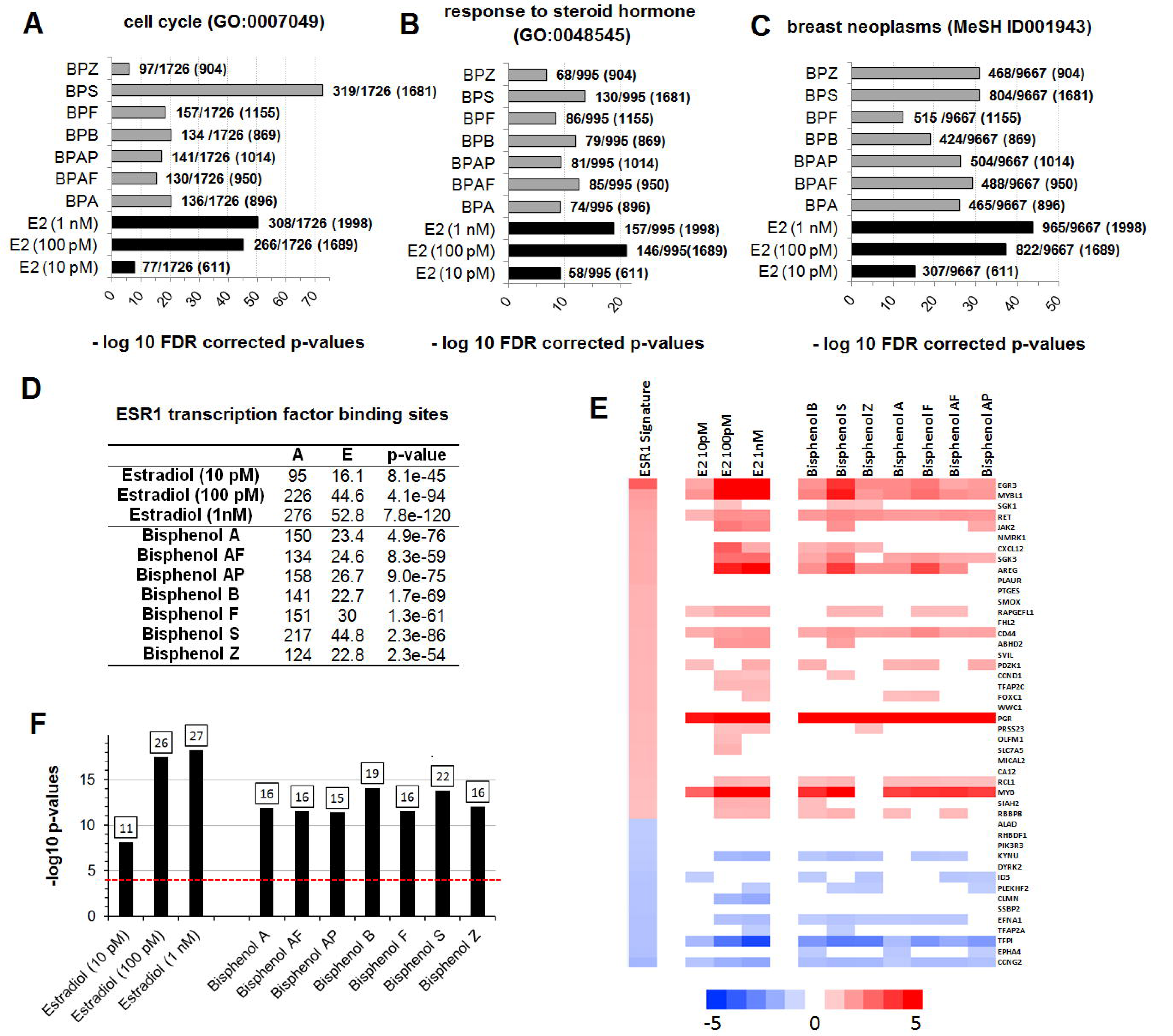
Alteration of MCF-7 transcriptome profiles caused by BPA alternatives bear the signature of ER activation. Transcriptome analysis of MCF-7 cells treated with BPA and alternative bisphenols reflect cell cycle changes **(A)**, response to hormone **(B)**, as well as an association to breast cancer **(C)**. The p-values are determined by hypergeometric calculation and adjusted using a Benjamini & Hochberg approach. Ratios show the total number of network objects belonging to each term in comparison to those disturbed by the treatments. The total of network objects (that is, all DEG) recognized by Metacore are indicated in parentheses **(D)**. Overrepresentation of ER binding motifs in the promoters of the differentially expressed genes. The analysis conducted with the transcription factor analysis tool of Metacore. A total of 1262 ER binding sites are found in the 42909 protein-based objects in the Metacore background list. A, number of targets in the activated dataset regulated by ESR1; E, expected mean of hypergeometric distribution; p-value calculated using hypergeometric distribution. **(E)** A gene expression biomarker confirms that BPA and bisphenol alternatives are ER agonists. Lists of statistically significant genes from MCF-7 cells treated with BPA and bisphenol alternatives or the natural hormone 17β-estradiol were examined against an ER gene expression biomarker signature consisting of 46 genes. The heat map shows the expression of genes in the biomarker after exposure to the indicated compound. Fold-change values for the ER biomarker are the average across 7 agonist treatments. **(F)** Bar plots showing the significance of the correlation by their −log10 p-values. Classification of activation or suppression required p<0.0001. The number of genes overlapping the ER biomarker is indicated at the top of each bar.

**Table 2.** List of the 5 most up- or down-regulated genes after treatment with BPA, BPAF or estradiol (E2). The transcriptome profiling was performed using the Affymetrix microarray platform.

| E2 (10 pM) |  | E2 (100 pM) |  | E2 (1 nM) |  | BPA |  | BPAF |  |
| --- | --- | --- | --- | --- | --- | --- | --- | --- | --- |
| ALDH1A3 | 0.4 | ALDH1A3 | 0.2 | ALDH1A3 | 0.2 | SEMA5A | 0.3 | ALDH1A3 | 0.3 |
| SEMA5A | 0.4 | PRICKLE2-AS3 | 0.2 | PSCA | 0.2 | ALDH1A3 | 0.3 | PSCA | 0.4 |
| PPARG | 0.5 | SEMA5A | 0.3 | TFPI | 0.2 | PSCA | 0.3 | RP11-706C16.7 | 0.4 |
| PSCA | 0.5 | PSCA | 0.3 | PRICKLE2-AS3 | 0.2 | PRICKLE2-AS3 | 0.3 | SEMA5A | 0.4 |
| IGFBPL1 | 0.5 | EPAS1 | 0.3 | EPAS1 | 0.3 | GABBR2 | 0.3 | EPAS1 | 0.4 |
| AC005534.8 | 2.6 | MYBL1 | 5.5 | AC106875.1 | 6.4 | AC106875.1 | 3.3 | AGR3 | 3.2 |
| MYB | 3.0 | AC106875.1 | 6.3 | MYBL1 | 6.5 | MYB | 3.7 | MYB-AS1 | 3.4 |
| AC106875.1 | 3.2 | GREB1 | 7.1 | GREB1 | 7.1 | GREB1 | 3.8 | MGP | 3.5 |
| GREB1 | 3.7 | PGR | 10.0 | PGR | 9.2 | MGP | 4.2 | MYB | 4.1 |
| PGR | 4.6 | MGP | 12.4 | MGP | 15.0 | PGR | 6.2 | PGR | 5.7 |
| BPB |  | BPF |  | BPS |  | BPZ |  | BPAP |  |
| ALDH1A3 | 0.3 | ALDH1A3 | 0.3 | ALDH1A3 | 0.2 | ALDH1A3 | 0.3 | ALDH1A3 | 0.3 |
| GABBR2 | 0.3 | SEMA5A | 0.3 | PSCA | 0.3 | TFPI | 0.3 | SEMA5A | 0.4 |
| PSCA | 0.3 | EPAS1 | 0.4 | SEMA5A | 0.3 | GABBR2 | 0.3 | GABBR2 | 0.4 |
| PRICKLE2-AS3 | 0.4 | PSCA | 0.4 | TFPI | 0.3 | PSCA | 0.4 | TFPI | 0.4 |
| EPAS1 | 0.4 | HCAR3 | 0.4 | PRICKLE2-AS3 | 0.3 | EPAS1 | 0.4 | EPAS1 | 0.4 |
| AGR3 | 3.3 | MYB | 3.9 | AC106875.1 | 5.4 | STC1 | 2.8 | STC1 | 3.1 |
| GREB1 | 3.8 | MYB-AS1 | 4.4 | MYB | 5.7 | ASCL1 | 2.9 | AL121578.2 | 3.1 |
| MYB | 4.1 | GREB1 | 4.5 | GREB1 | 6.0 | AGR3 | 3.3 | MYB | 3.8 |
| MGP | 4.4 | MGP | 5.0 | MGP | 7.8 | MYB-AS1 | 3.4 | AGR3 | 3.8 |
| PGR | 6.8 | PGR | 6.5 | PGR | 9.5 | PGR | 5.8 | PGR | 5.8 |

Statistical values derived from gene ontology analysis can have limited reliability due to the background set including only the genes that are likely to be expressed in the experiment (Tipney and Hunter, 2010). Genes disturbed by chance have a higher probability to be associated with endocrine ontologies in an endocrine sensitive tissue such as the breast. This could result in an increased false positive rate. In order to circumvent this problem, we applied a biomarker approach to predict ER modulation. The gene expression biomarker consists of 46 genes which exhibit consistent expression patterns after exposure to 7 agonists and three antagonists of ER determined using MCF-7 microarray data derived from the CMAP 2.0 dataset (Ryan et al., 2016). This gene expression biomarker has a balanced accuracy for prediction of ER activation of 94%. Gene expression profiles from the cells treated with BPA and BP alternatives were compared to the ER biomarker using the Running Fisher algorithm as previously described (Ryan et al., 2016). The cut-off value for statistical significance was p-value ≤ 0.0001 after a Benjamini Hochberg correction of α = 0.001. All transcriptome profile alterations resulting from exposure to BPA and all 6 BP alternatives (Figure 3E) were highly statistically significant resulting from expression of biomarker genes in a pattern highly similar to that of the biomarker (Figure 3F).

In order to confirm estrogenic effects provoked by BPA and BP alternatives, RNAs extracted from MCF-7 cells treated with BPAF (0.08 μM), BPA (0.36 μM) and estradiol (1 nM) were subjected to total transcriptome RNA sequencing (RNA-seq) analysis using the Illumina HiSeq 2500 system. This also allowed us to compare the sensitivity of RNA-seq and microarrays to determine estrogenic effects. We have performed pairwise comparisons to determine the list of DEGs following the same criteria used for the microarray analysis. Only 12-21% of the DEGs identified by RNA-seq were also found to be altered on the microarrays (Figure 4B). The fold changes of these genes found commonly altered in both the microarray and the RNA-seq analysis in comparison to their respective controls were well correlated (Figure 4A) for 17β-estradiol (Pearson r = 0.81), BPA (r = 0.86) and BPAF (r = 0.81). Overall, the RNA-seq method was more sensitive and identified 2-3 times more significantly altered genes compared to the microarray method (Figure 4B). A total of 5091, 2930 and 3093 genes were significantly altered by estradiol, BPA and BPAF, respectively. Genes commonly found altered after the application of both transcriptome profiling methods generally had a higher fold change by the RNA-seq platform (Supplementary File 3). The ontology analysis based on RNA-seq data gave similar results to our previous calculations based on the microarray data and confirms estrogenic effects of BPA and BPAF (Figure 4C). Although more DEGs related to cell cycle function were found by RNA seq than by microarray, the - log10 p-value for BPA was lower because the total number of DEGs found by RNA seq was higher than by microarray. We similarly applied the ER gene expression biomarker to detect ER agonists after analysis with the Illumina sequencing platform (Figure 4D and 4E). The RNA-seq platform gave results similar to that of Affymetrix microarrays confirming estrogenic effects of BPA alternatives.

**Figure 4.**
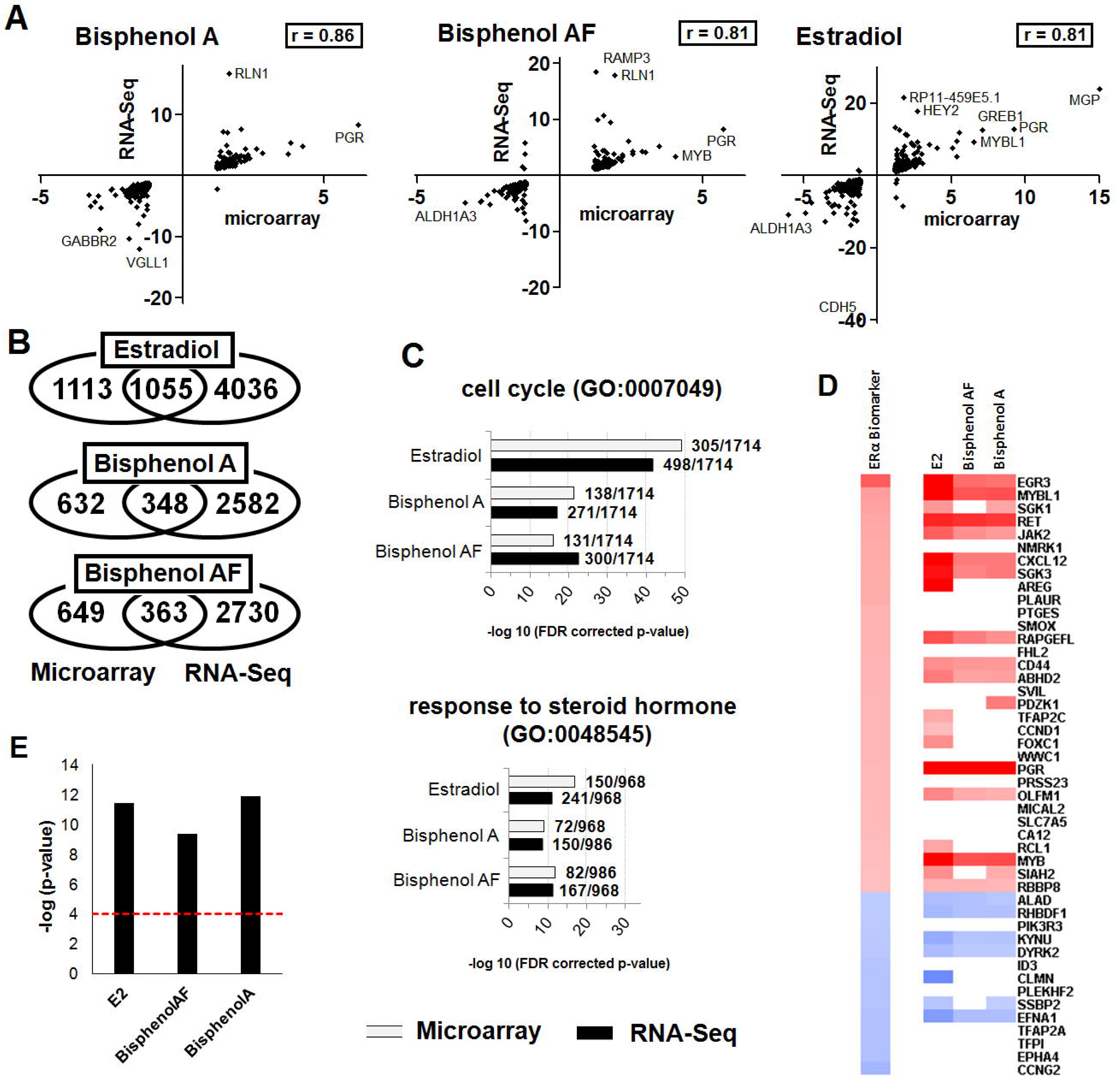
Comparison of RNA-seq and microarray platforms in determining endocrine disrupting effects of BPA and bisphenol alternatives. RNA extracted from MCF-7 cells was subjected to a full transcriptome profiling using the Illumina RNA sequencing or the microarray technology under similar conditions. **(A)** Pearson correlation coefficients between the RNA-seq and microarray data. The fold changes in gene function having an altered expression by the two methods are presented. **(B)** Venn diagrams showing the number of genes uniquely or commonly disturbed. **(C)** Gene ontology analysis of terms associated with MCF-7 hormone induced proliferation in the transcriptome profiles obtained by RNA-seq or microarray analysis. Ratios are showing the total number of network objects belonging to each term in comparison to those disturbed by the treatments. The total of network objects (that is, all DEG) recognized by Metacore are indicated in parentheses. **(D)** Heat map of genes whose expression was statistically significantly altered examined against an ER gene expression biomarker. **(E)** Bar plots showing the significance of the correlation by their - log10 p-values.

## Discussion

Although human populations are exposed to a wide range of BPA alternatives, their estrogenic effects have never been thoroughly determined. We present here the first comprehensive, side-by-side comparison of the estrogenic effects of BPA and 6 BP alternatives (BPZ, BPAF, BPAP, BPB, BPF, BPS) found in foodstuffs and human fluids using both cell proliferation and gene expression profile assays. Our study revealed that the 6 bisphenols introduced in “BPA-free” plastics displayed estrogenic effects with BPAF, BPB and BPZ being more potent than BPA, although their estrogenic potential remained in the same range as BPA. The comparison of transcriptome profiles revealed a signature demonstrating that these effects were mediated via ER (Figure 3). Our results highlight the need to conduct studies to examine possible pathological effects in laboratory animals at concentrations relevant for human exposures. Similar transcriptome alterations should also be confirmed in other cell lines or primary cells. In addition, combined effects of these bisphenols should be explored in further studies to determine the relevance of these estrogenic effects in terms of human health risk assessment. In particular, our results call for the relevance of human exposure to BPA alternatives in hormone-dependent breast cancer progression to be investigated.

The structure of BPA alternatives (Figure 1) provide insights into the possible mechanisms of action toward ER. BPA binding to ER is mainly driven by van der Waals' forces and hydrogen bond interactions (Li *et al.,* 2015). BPA and 17β-estradiol bind in a similar manner, with two phenol rings pointing to the two ends of the ER hydrophobic pocket. Differences in estrogenic activities between the different BPA alternatives may be due to the different groups present at the bridge between the two phenol rings. Their hydrophobicity is driving their affinity, since the matching of the group carried by the methylene bridge is known to determine binding affinity towards the hydrophobic surface of the ER binding site (Endo *et al.,* 2005). Such a mechanism of ER interaction is supported by the hydrophobicity ranking of bisphenols (logP values BPZ > BPAF > BPAP > BPB > BPA > BPF > BPS), which closely corresponds to their AC50 toxicity score.

Future studies need to explore effects at lower doses, since EDCs can elicit a nonmonotonic response that significantly deviates from the usual sigmoidal curve (Vandenberg *et al.,* 2012). In another transcriptomics study in which MCF-7 cells were exposed to varying concentrations of BPA, a weak gene activation peak at a very low concentration range (~0.1 nM) was observed in addition to the main peak of gene activation (Shioda et al., 2013). Additionally, an investigation of the combinatorial effects of bisphenols would reflect real-world exposure conditions even if estrogenic effects of chemical mixtures *in vitro* are predicted by the use of concentration addition models (Evans *et al.,* 2012).

Our data allows a direct comparison of the sensitivity of RNA-seq and microarray hybridisation in the determination of the transcriptome signature of endocrine disrupting effects. Overall, the application of RNA-seq resulted in the detection of more statistically significant alterations in gene expression than the microarray method. These genes generally had a higher fold change leading to p-values associated with gene expression biomarkers being lower. Overall, RNA sequencing appears to be more sensitive than microarray analysis. This was also the conclusion of studies performed in other cell lines (Perkins *et al.,* 2014), mesenchymal stem cells (Li *et al.,* 2016) and rat tissues (Perkins et al., 2014). In a study of 498 primary neuroblastomas, RNA-seq outperformed microarrays in determining the transcriptomic characteristics of these cancers, while RNA-seq and microarray performed similarly in clinical endpoint prediction (Zhang *et al.,* 2015). Both technologies are suitable to detect estrogenic effects of BPA alternatives using the gene expression biomarker of ERα activation. Although the genes weakly expressed were differently detected by the two platforms, the biomarker signatures were comparable. However, it is perhaps noteworthy that the microarray processing and analysis pipelines are better standardized and could thus be more reliable in comparing experiments performed by different laboratories. Further intra- and inter-experimental comparisons would be necessary to conclude which method is the most sensitive and reliable to determine transcriptome signatures.

The mechanisms of action resulting in toxic effects from BPA at low levels of exposure have long remained elusive due to its relatively low affinity to ER. Higher affinities in the circulating exposure range for BPA have been reported on membrane estrogen receptor GPR30 / GPER at a concentration of 1 nM (Sheng et al., 2012). BPA is very potent at inducing rapid non-genomic responses from membrane surface receptors at low concentrations (1-100 pM) (Wozniak *et al.,* 2005). For example, BPA administration inhibited meiotic maturation of Zebrafish oocytes through a G protein-coupled ER-dependent epidermal growth factor receptor pathway (Fitzgerald *et al.,* 2015). BPA can also induce adipogenesis through other receptors such as peroxisome proliferator-activated receptor gamma (PPARγ) (Ahmed and Atlas, 2016). Surprisingly, this latest study provided evidence that BPS is a more potent adipogen than BPA in inducing 3T3-L1 adipocyte differentiation and lipid accumulation. Overall, effects of BPA alternatives have been poorly investigated. Recent studies have revealed that less studied endocrine-related systems such as glucocorticoid, PPAR, monoamine, noradrenaline and serotonin pathways could be targets of endocrine disruption (Filer *et al.,* 2014).

In adult humans, BPA is considered to be rapidly metabolized by the liver, with elimination virtually complete within 24 h of exposure, although chronic exposures could result in accumulation if BPA distributes within tissues that slowly release BPA (Stahlhut *et al.,* 2009). In an investigation of concentrations of BPA metabolites in the urine of 112 pregnant women (ethnically and racially diverse), total BPA consisted of 71% BPA glucuronide, 15% BPA sulfate and 14% unconjugated BPA (Gerona et al., 2016). These metabolites have long been believed to be biologically inactive because they are not thought to be ER agonists. However, this does not preclude effects on other receptor systems or the creation of equilibrium between the metabolite and the parent if the metabolite is long-lived. For instance, a recent study showed that BPA glucuronide can induce lipid accumulation and differentiation of pre-adipocytes (Boucher *et al.,* 2015). Metabolites of the main BPA alternatives have not been routinely measured in human biomonitoring studies, which may greatly underestimate real world exposures. Moreover, BPA metabolites may be deconjugated by β-glucuronidase, which is particularly active in the placenta and fetal liver resulting in increased fetal exposure (Ginsberg and Rice, 2009). Overall, the assessment of potential health effects arising from exposure to bisphenols would be greatly facilitated if biomonitoring studies examining reference populations like those of the U.S. CDC’s NHANES programme, were to begin routine examination of human fluids for BPA alternatives as well as their metabolites.

Even though disruption of ER function is one of the most investigated endocrine-related targets in the ToxCast data (18 assays) (Judson et al., 2015), our analysis of the high throughput system (HTS) employed has revealed discrepancies between our results and those of this screen with some of these ToxCast assays failing to detect estrogenic effects (Table 1). This was especially the case for BPF which was found negative in 9 out of the 11 assays scrutinized. The two systems of analysis detecting the estrogenic capability of BPF were the Attagene assays. BPF was found to be less potent in these assays than in the present study (Table 1). Our results are in agreement with previously published estrogenic effects of BPF reporting an EC50 of 0.82 μM in a ER luciferase assay (Rosenmai et al., 2014). BPAP on the other hand was found to be more potent in the ToxCast assays, although the difference within the 2 current assays was minimal and could be explained by different grades of chemical purity. Another reason for these differences could be the different sensitivities of HTS assays. These are frequently based on heterologous expression of reporter genes in non-hormone-responsive cell lines that possibly do not possess all the co-factors necessary to transduce the hormone signal, thus resulting in a decreased sensitivity to detect relevant signals.

The US EPA has proposed to substitute the Endocrine Disruptor Screening Program Tier 1 assays with in vitro data in order to reduce the cost and time of toxicity testing, as well as animal use (Browne *et al.,* 2015). One way forward to comply with these objectives would be to integrate gene expression profiling into the HTS system. Transcriptome profiles resulting from exposure to a given chemical could be correlated to signatures of a wide range of chemicals using the Library of Integrated Network-based Cellular Signatures (LINCS) database, which contains signatures from around 4000 chemicals screened in approximately 17 cell lines (http://www.lincsproject.org). This could be used as a “Tier 0” assay to further prioritize targeted *in vitro* testing in the context of toxicity testing programmes (Ryan et al., 2016).

## Conclusion

We have detected estrogenic effects of BPA alternatives by the application of a gene expression biomarker of ERα activation, whose results are corroborated by functional cellular assays. Our comparison of microarray and RNA-Seq technologies showed that both platforms are suitable for the use of this ERα gene expression biomarker.

Recently, the plastics manufacturing industry have turned to alternative bisphenols to produce their “BPA-free” products, often with little toxicology testing. Our data show that some of these BPA alternatives are even more potent ER activators than BPA, suggesting that “bisphenol-free” labels on consumer products may need to be more informative.

## Acknowledgements

This work was funded by the Sustainable Food Alliance (USA) and Breast Cancer UK, whose support is gratefully acknowledged. This study has been subjected to review by the U.S. EPA National Health and Environmental Effects Research Laboratory and approved for publication. Approval does not signify that the contents reflect the views of the Agency, nor does mention of trade names or commercial products constitute endorsement or recommendation for use. We thank Drs. Richard Judson and Charles Wood for critical review of the manuscript and Dr. John Rooney for technical assistance.

## Supplementary files

### Supplementary File 1 Volcano plots of the MCF7 transcriptome profile changes induced by BPA and bisphenol alternatives

Alterations in MCF7 transcriptome profiles induced by a 48h exposure to three concentrations of 17β-estradiol and the AC(50) of 7 bisphenols were determined using the Affymetrix GeneChip^®^ Human Transcriptome Array 2.0. The volcano plots show the log 2 fold changes and the −log10 p-values in transcript cluster expression compared to the negative control. Data selected at the cut off values p < 0.05 and fold change > 1.2 for the functional analysis were represented by red dots.

### Supplementary File 2 Alterations of the MCF7 transcriptome induced by BPA and bisphenol alternatives

MCF-7 cells were subjected to a full transcriptome profiling using the Affymetrix microarray platform. The list of genes having their expression altered by the treatment with the BPA alternatives or three concentrations of 17β-estradiol is provided with their fold-changes and the p-values. Data were selected at the cut off values p < 0.05 and fold change > 1.2.

### Supplementary File 3 Alterations of the MCF7 transcriptome induced by BPA, BPAF, and 17β-estradiol

MCF-7 cells were subjected to a full transcriptome profiling using the Illumina RNA sequencing technology. The list of genes having their expression altered by the treatment with BPA, BPAF, or 17β-estradiol is provided with their fold-changes and the p-values. Data were selected at the cut off values p < 0.05 and fold change > 1.2

